# Microtubule-associated proteins MAP7 and MAP7D1 promote DNA double strand break repair in the G1 cell cycle phase

**DOI:** 10.1101/2023.01.17.524354

**Authors:** Arlinda Dullovi, Meryem Ozgencil, Vinothini Rajvee, Wai Yiu Tse, Pedro R Cutillas, Sarah A Martin, Zuzana Hořejší

## Abstract

The DNA-damage response is a complex signalling network that guards genomic integrity. The microtubule cytoskeleton is involved in the repair of DNA double-strand breaks; however, little is known about which cytoskeleton-related proteins are involved in DNA repair and how. Using quantitative proteomics, we discovered that microtubule associated proteins MAP7 and MAP7D1 interact with several DNA repair proteins including DNA double-strand break repair proteins RAD50, BRCA1 and 53BP1. We observed that downregulation of MAP7 and MAP7D1 leads to increased phosphorylation of p53 after γ- irradiation. Moreover, we determined that the downregulation of MAP7D1 leads to a strong G1 arrest and that the downregulation of MAP7 and MAP7D1 in cells arrested in G1 negatively affects DNA repair, recruitment of RAD50 to chromatin and localisation of 53BP1 to the sites of damage. These findings describe for the first time a novel function of MAP7 and MAP7D1 in cell cycle regulation and cellular response to DNA double-strand breaks.

## Introduction

The DNA-damage response (DDR) is a complex signalling network that protects cells against physical and chemical lesions, thus preserving their genomic integrity^1^. Genetic aberrations in DDR components lead to genome instability and a number of diseases including cancer^2–4^. Although the majority of DDR occurs in the nucleus, there is an increasing evidence that cytoplasmic processes such as sensing of double stranded DNA inside ruptured micronuclei by cGAS/STING innate immunity pathway^5^, autophagy^6^, translational processes and energy metabolism^7–9^ are also crucial for DNA repair and cellular homeostasis after DNA damage.

Microtubules are functionally and structurally important components of the eukaryotic cytoskeleton, formed from highly conserved α/β-tubulin heterodimers. They participate in the majority of the cellular processes, including trafficking of organelles and proteins, cell motility and cell division^10^. Microtubules have been implicated in the repair of the DNA double-strand breaks (DSBs) by affecting the directional mobility of damaged DNA via the LINC complex and by dynamic nuclear microtubule filaments in asynchronous cells or by the induction of microtubule dynamic stress response in G0 and G1 cell cycle phases^11–13^. Moreover, KIF18B and KIF2C, two kinesins which act as motor proteins on microtubules, are involved in DSB repair^14, 15^. Nevertheless, the scale of processes through which microtubules affect DDR, the number of cytoskeleton-related proteins involved in DDR and the exact molecular mechanisms behind these processes remain largely unknown.

The structure of the microtubule cytoskeleton is regulated by the interactions between microtubules with numerous microtubule-associated proteins (MAPs). MAPs control microtubule polarization, depolarisation and stabilisation, connect microtubules to the membranes and regulate microtubule motor motility and cell migration^16^. MAP7 is a conserved, non-motor MAP protein that participates in the regulation of microtubule organisation, assembly dynamics and stability^17, 18^. It recruits kinesin-1 to the microtubules to direct organelle and protein trafficking from the nucleus to the cellular periphery^19–26^. MAP7 and its paralogue MAP7D1 regulate the migration and invasion of gastric cancer cells^27^ and the migration, adhesion and cell cycle progression in cervical cancer cells through NF-αB and Wnt5α signalling^28–30^.

In this study, we show that MAP7 and MAP7D1 interact with several DNA repair proteins including RAD50, BRCA1 and 53BP1. Moreover, we determined that the downregulation of MAP7D1 leads to strong G1 arrest and that downregulation of MAP7 and MAP7D1 in cells arrested in G1 negatively affect DSB repair, RAD50 recruitment to chromatin and localisation of 53BP1 to the site of damage. Taken together, these findings describe for the first time an important function of MAP7 and MAP7D1 in cell cycle regulation and cellular response to DNA damage caused by γ-irradiation.

## Results

### The DDR proteins BRCA1, RAD50, MLH1 and XPC interact with microtubule associated proteins MAP7 and MAP7D1

Phosphorylation is a highly important posttranslational modification in DNA damage signalling. Previously, we and others have used peptide-pull down assays as a direct means of attributing specific binding functions to known sites phosphorylated by CK2 in DDR proteins ^31, 32^. We have now applied this approach to identify proteins binding to four *in vivo* phosphorylated sites from DDR proteins RAD50 (Thr690), BRCA1 (Ser1336), MLH1 (Ser477) and XPC (Ser883, Ser884). These sites are conserved, reside within consensus sites for CK2, share similar sequence features (Figure 1A), are mutated in cancer and/or their phosphorylation is known to affect (or is affected) by the DDR ^33–35^. Thus, we reasoned that these putative CK2 sites may be of critical functional importance and that their analysis might provide further insight into regulation of the DDR. We designed biotinylated 20 amino acid (aa) peptides encompassing the specific phosphorylated amino acids of BRCA1, RAD50, MLH1 and XPC that were either non- phosphorylated or phosphorylated. The biotinylated peptides were coupled to streptavidin beads and used to pull down proteins from HeLa nuclear extracts. Quantitative proteomic analysis revealed that all four sites bind proteins in a phospho-dependent manner (Figure 1D). Each of the phosphorylated sites bound various proteins including those implicated in DDR and cell cycle regulation as well as proteins involved in other cellular processes; some of these proteins have already known phospho-binding domains, while others do not (Figure 1B). Interestingly, the microtubule associated protein MAP7 featured prominently in all of the phosphorylated RAD50, BRCA1, MLH1 and XPC peptide pull-downs and the MAP7 paralogue MAP7D1 also featured in the phosphorylated BRCA1 peptide pull-down. The binding of MAP7 by the RAD50 phosphorylated peptide was further confirmed by western blotting (Figure 1C). These results suggested that MAP7 (and possibly MAP7D1) may act as a general interaction scaffold that connects the DDR and cytoskeleton machinery.

**Figure 1.**
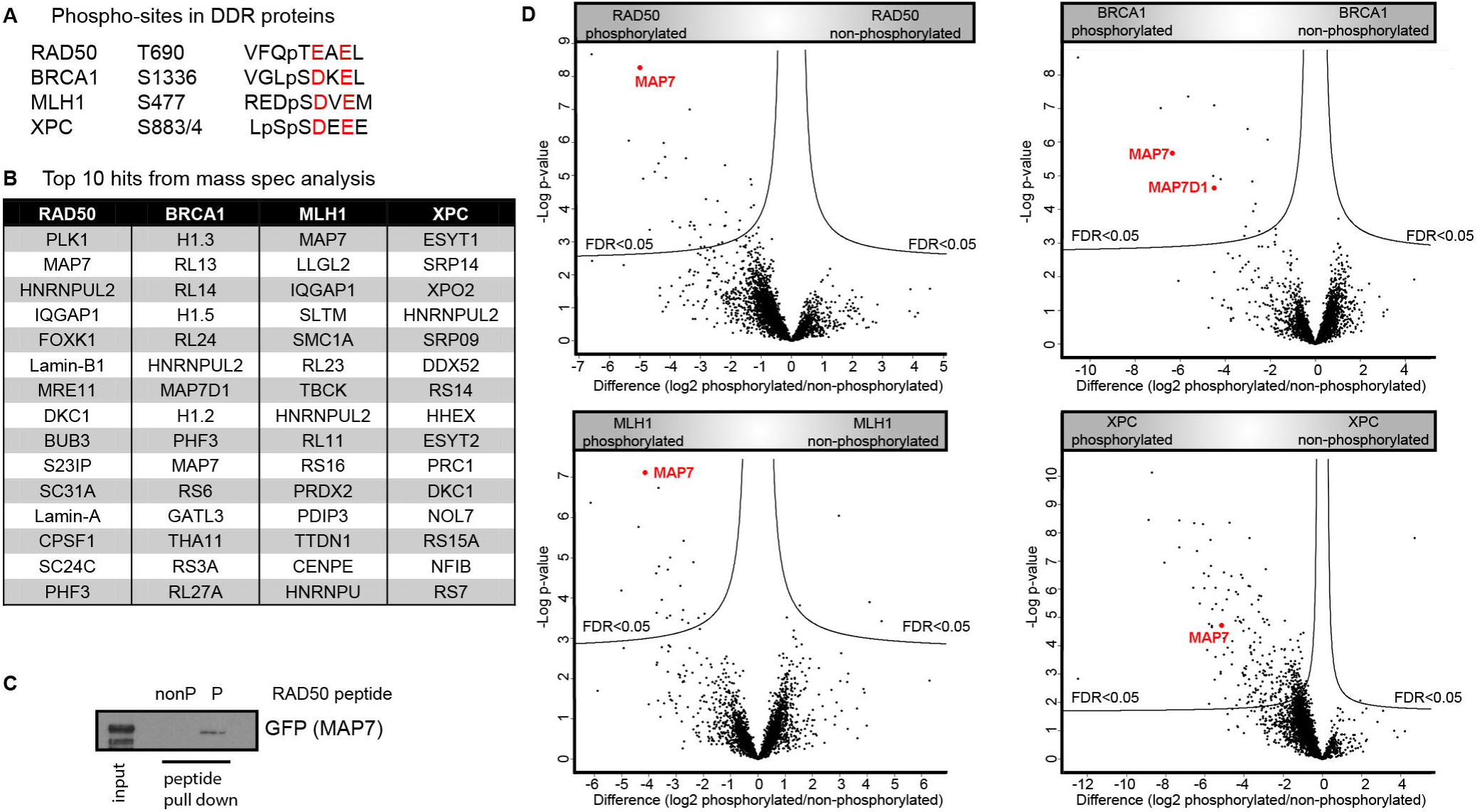
Phosphorylated peptides from RAD50, BRCA1, MLH1 and XPC interact with MAP7. (A) Peptides from four DDR proteins containing the similar feature S/TE/DXE. Amino acids with a negative charge are highlighted in red. (B) Top 10 hits from the proteomics of peptide pulldowns from HeLa nuclear extract performed with pairs of phosphorylated and non-phosphorylated peptides from BRCA1, RAD50, MLH1 and XPC. (C) Western blot analysis of peptide pull-down from HEK293T whole cell extract overexpressing GFP-MAP7 with non-phosphorylated (nonP) and phosphorylated (P) RAD50 peptides.(D) Volcano plots compare proteins binding to phosphorylated and non-phosphorylated peptides. A Benjamini-Hochberg cut-off of 5% with an s0 of (0.1) was used to select significant binders.

Therefore, we sought to confirm these interactions within the context of full-length MAP7 and MAP7D1 proteins. Western blotting of immuno-precipitates from HEK293T whole cell extracts confirmed the interactions between the full-length endogenous MAP7 and MAP7D1 with RAD50 and BRCA1 (Figure 2A, B) and between GFP-tagged full-length MAP7 with endogenous MLH1 and XPC (Figure S2A, B). These results indicate that MAP7 and MAP7D1 form a complex with several DDR proteins under physiological conditions.

**Figure 2.**
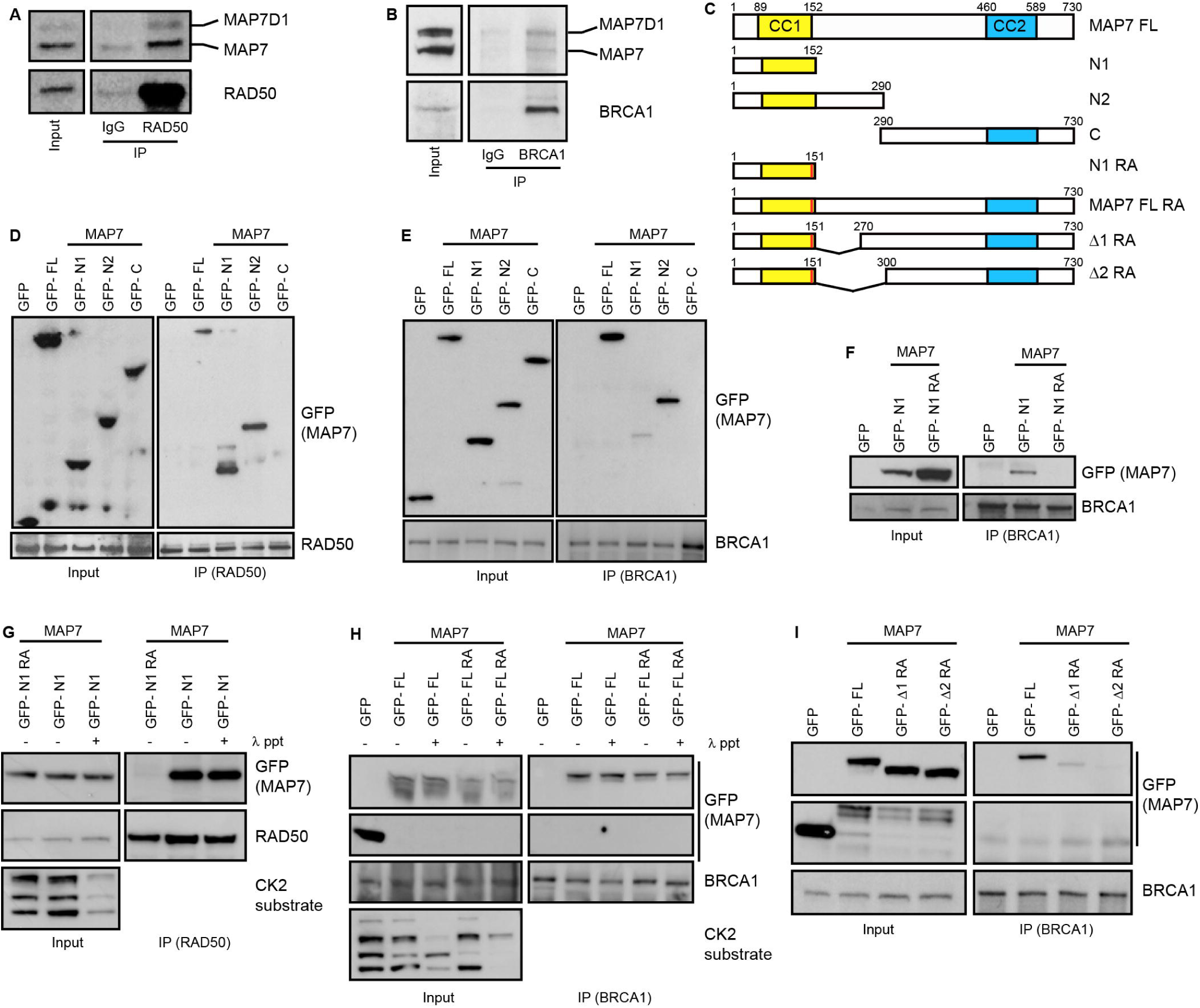
Interactions between MAP7 and MAP7D1 with RAD50 and BRCA1. (A) Immuno-precipitation of endogenous MAP7 and MAP7D1 with RAD50 from HEK293T whole cell extract (both MAP7 and MAP7D1 are recognised by MAP7 antibody). (B) Immuno-precipitation of endogenous MAP7 and MAP7D1 with BRCA1 from HEK293T whole cell extract. (C) Schematic representation of full-length MAP7 and MAP7 constructs used in immuno-precipitations. CC1 and CC2 are two coil-coiled domains. (D) Domain mapping by immuno-precipitation of full-length GFP-MAP7 (GFP-FL) and MAP7 truncation constructs by endogenous RAD50 from HEK293T extract overexpressing GFP-MAP7 constructs. (E) Domain mapping by immuno-precipitation of full-length GFP-MAP7 (GFP- FL) and GFP-MAP7 truncation constructs by endogenous BRCA1 from HEK293T extract overexpressing GFP-MAP7 constructs. (F) Immuno-precipitation of GFP-tagged MAP7 N- terminal fragment (aa1-152) wild type (GFP-N1) or with Arg144 and 145 mutated to Ala (GFP-N1RA) with endogenous BRCA1 from HEK293T extract overexpressing GFP-MAP7 constructs. (G) Immuno-precipitation of GFP-N1 with endogenous RAD50 from HEK293T whole cell extract overexpressing GFP-MAP7 construct, untreated or treated with λ- phosphatase. (H) Immuno-precipitation of full length MAP7 (GFP-FL) wild type and MAP7 mutated on arginine 144/145 to alanine (GFP-FL RA) with endogenous BRCA1 from HEK293T whole cell extract treated or untreated with λ-phosphatase. (I) Further domain mapping using MAP7 constructs mutated on Arg144/145 to Ala with deletion between aa151-270 (GFP- Δ 1 RA) and 151-300 (GFP- Δ 2 RA).

To determine which domains of MAP7 interact with the DDR proteins, we expressed various truncation constructs of GFP-tagged MAP7 in HEK293T cells (Figure 2C). MAP7 constructs with truncated C-terminus (aa1-152 and aa1-290) were able to bind all four DDR proteins, while an N-terminal truncation of MAP7 (aa290-730) abolished these interactions (Figure 2D, E; Figure S2A, B). We continued to identify specific MAP7 residues that are required for the interactions by mutagenesis of the N-terminal part of MAP7 using immuno-precipitation. By mutating positively charged amino acids, which may be responsible for the phospho-specific interaction, we have discovered that the mutation of two conserved Arg residues (Arg144 and Arg145) to Ala disrupts the interaction of the shorter N-terminal MAP7 fragment (aa1 to 152) with BRCA1 and RAD50 (Figure 2F, G). However, the interaction of this N-terminal fragment with RAD50 was not abolished after treatment of the whole cell extract with λ-phosphatase (Figure 2G), indicating that these two Arg residues do not interact with the phosphorylated Thr690 of RAD50. In addition, the λ-phosphatase treatment of the whole cell extract from the HEK293T cells expressing either full length MAP7 wt or the double Arg to Ala mutated MAP7, did not alter the BRCA1 interaction with MAP7 by immuno-precipitation (Figure 2H). By creating further deletion mutations of MAP7, we have determined that the deletion of the MAP7 region between aa152-300 in combination with the double Arg to Ala mutation abolishes MAP7 interaction with BRCA1 and RAD50 (Figure 2I, 2SC). These data raised the possibility that the interaction between MAP7 and the DDR proteins may be direct but requires multiple binding sites, several of which are not dependent on phosphorylation. Alternatively, the interaction is indirectly bridged by other protein(s) binding to the MAP7 region that spans between aa144 to aa300.

Next, we tested if a similar region within MAP7D1 is also responsible for the interactions with DDR proteins. We deleted aa216-424, which correspond to aa144-300 within MAP7 (Figure S2D). This deletion of mCherry-tagged MAP7D1 resulted in a slightly reduced the interaction with endogenous BRCA1 but did not affect the interaction with endogenous RAD50 (Figure S2E, F). This may be because only part of the deleted region is conserved between MAP7 and MAP7D1, while both proteins may require different parts of their non-conserved regions for the interactions with the DDR proteins (Fig. S2C).

### Direct interaction between MAP7 and BRCA1 is phosphorylation-dependent and regulated by CK2 kinase

To determine if the interaction between MAP7 and the DDR proteins is direct, we purified full-length Flag-tagged RAD50 wild type (Flag-RAD50 wt) and Flag-tagged RAD50 mutated on Thr690 to Ala (Flag-RAD50 TA) using high-salt (4M) conditions. The purified proteins were used in *in vitro* binding experiments with GFP-MAP7 immobilized on GFP-trap beads washed with high-salt (4M) buffer. RAD50 wt efficiently bound to the GFP- MAP7, whereas RAD50 TA exhibited severely reduced binding to the GFP-MAP7 (Figure 3A). Likewise, λ-phosphatase treatment reduced RAD50 binding to GFP-MAP7 (Figure 3B). Similar experiments demonstrated that purified Flag-tagged MAP7 interacts with purified GFP-BRCA1 wt but this interaction is significantly lower with GFP-tagged BRCA1 mutated on Ser1336 to Ala (GFP-BRCA1 SA) or when GFP-BRCA1 wt is dephosphorylated by λ-phosphatase (Figure 3C). These data indicate that MAP7 binds to RAD50 and BRCA1 directly and that the direct interaction is dependent on phosphorylated RAD50 Thr690 and on phosphorylated BRCA1 Ser1336. Furthermore, these and the previously described data suggest that the interaction between MAP7 and the DDR proteins is also bridged by some other factors and that this indirect interaction is independent on phosphorylation.

**Figure 3.**
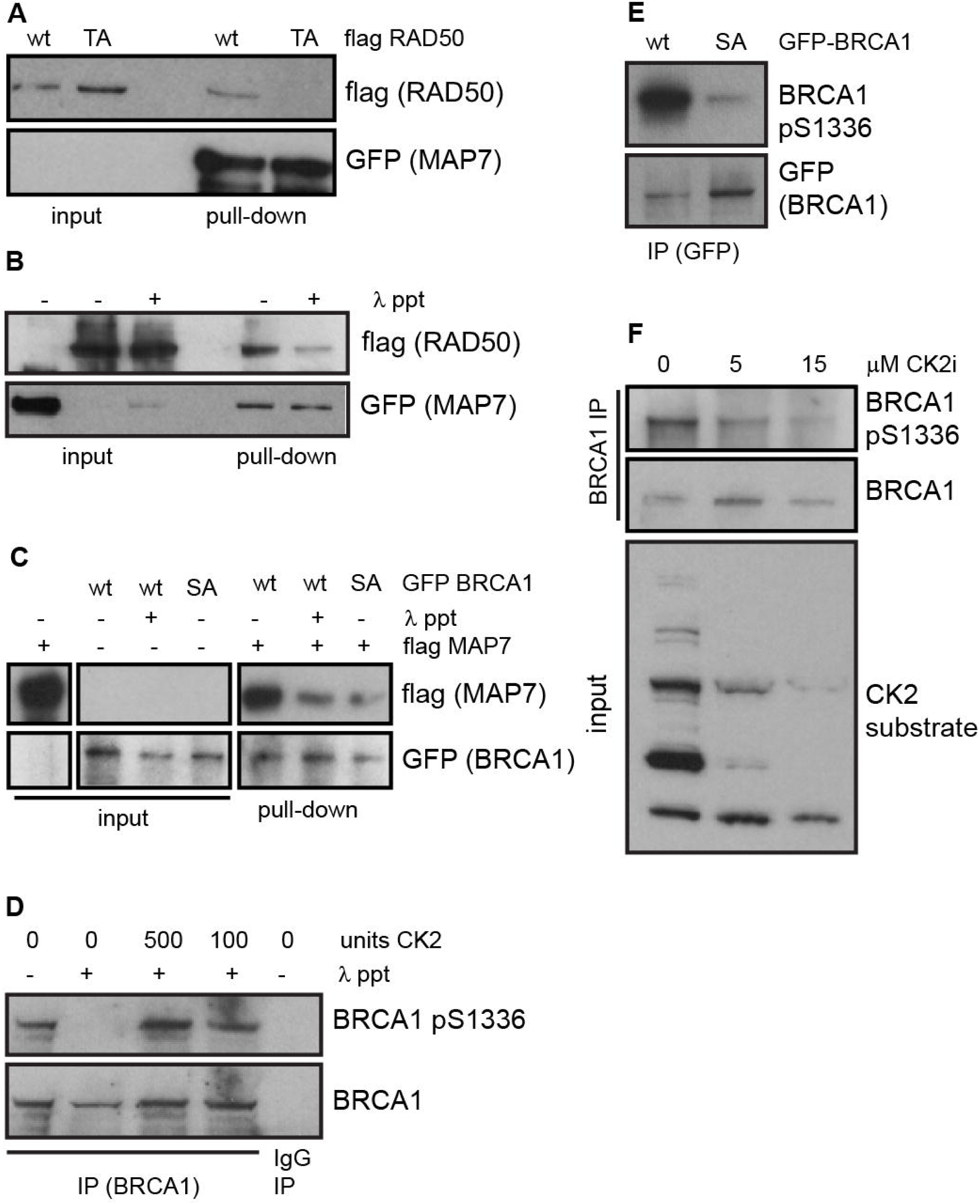
Direct interactions between MAP7, BRCA1 and RAD50 are phosphorylation-dependent and regulated by CK2. (A) Pull down of Flag-RAD50 wt or TA with GFP-MAP7. Proteins were purified from HEK293T whole cell extracts by immuno-precipitation followed by high salt (4M NaCl) washes. Flag-tagged proteins were eluted from beads using Flag peptide. (B) Pull down of Flag-RAD50 wt untreated or treated with λ-phosphatase with GFP-MAP7. Proteins were purified from HEK293T whole cell extracts by immuno-precipitation followed by high salt (4M NaCl) washes. Flag-tagged proteins were eluted from beads using Flag peptide. (C) Pull down of Flag-tagged MAP7 with GFP-BRCA1 wt untreated or treated with λ- phosphatase or GFP-BRCA1 SA mutation. Proteins were purified from HEK293T whole cell extracts by immuno-precipitation followed by high salt washes (4M NaCl). Flag-tagged proteins were eluted from beads using Flag peptide. (D) Immuno-precipitation of endogenous BRCA1 from HEK293T whole cell extracts. Proteins on beads were untreated or treated with λ-phosphatase followed by re-phosphorylation by recombinant CK2. (E) Immuno-precipitation of GFP-BRCA1 wt and GFP-BRCA1 SA from HEK293T whole cell extract. Immuno-precipitation of endogenous BRCA1 from HEK293T whole cell extracts. Proteins on beads were untreated or treated with λ-phosphatase followed by re-phosphorylation by recombinant CK2. (F) Immuno-precipitation of endogenous BRCA1 from HEK293T whole cell extract untreated or treated with two different doses of 5µM or 15µ CK2 inhibitor.

Since our results suggest that the phosphorylation of BRCA1 and RAD50 plays some role in their direct interaction with MAP7, we investigated which kinase is responsible for their phosphorylation. The peptide sequences of the four DDR proteinsthat pulled down MAP7 in our initial experiments contain the primary consensus target site for the serine/threonine protein kinase CK2. While MLH1 Ser 447 has been previously described as a CK2 site^35^, the kinase phosphorylating the other three sites has not yet been identified. We therefore raised a rabbit polyclonal antibody against a synthetic peptide phosphorylated on BRCA1 serine 1336, which was able to recognize immuno-precipitated GFP-BRCA1 wt overexpressed in HEK293T cells, while it did not detect overexpressed GFP-BRCA1 SA (Figure 3E). To prove the phospho-binding specificity of the antibody, we immuno- precipitated endogenous BRCA1. An aliquot of the beads binding BRCA1 was treated with λ-phosphatase and subsequently re-phosphorylated by recombinant CK2. The antibody failed to recognize dephosphorylated BRCA1 on western blot but readily detected a band corresponding to endogenous BRCA1 in the samples untreated with λ-phosphatase and samples that were re-phosphorylated by CK2 (Figure 3D). These data indicate that CK2 can phosphorylate BRCA1 Ser1336 *in vitro*. In addition, while the phospho-specific antibody recognised endogenous BRCA1 immuno-precipitated from the whole cell extract on western-blot, the levels of phosphorylated BRCA1 from whole cell extracts treated with a CK2 inhibitor were much lower (Figure 3F), suggesting that BRCA1 Ser1336 is also phosphorylated by CK2 *in vivo*.

### The interaction between MAP7 and the DDR proteins is not altered by nocodazole, paclitaxel and DNA damage treatment

To determine in which cellular compartment MAP7 interacts with the DDR proteins, we performed a proximity ligation assay in U2OS cells transfected with Flag-tagged MAP7 using Flag, RAD50 and BRCA1 antibodies. The results indicate that MAP7 interacts with RAD50 and BRCA1 both in the cytoplasm and the nucleus. However, in 30% or 50% of cells, MAP7 interacts with RAD50 and BRCA1 (respectively) predominantly in the nucleus (Figure 4A, B). Next, we investigated whether the interactions between MAP7/MAP7D1 and the DDR proteins may be mediated by microtubules. To this end, we treated HEK293T cells with nocodazole (an inhibitor causing microtubule disassembly) or paclitaxel (stabilizes the microtubule polymer and protects it from disassembly). Neither treatment affected the interaction between endogenous BRCA1 or RAD50 and GFP-MAP7 (Figure 4C-F). In accordance, nocodazole and paclitaxel treatment also had no effect on the interaction between mCherry-MAP7D1 and endogenous BRCA1 (Figure 4G, H), indicating that the interactions are not bridged by microtubules. Similarly, treatment of cells overexpressing GFP-MAP7 with γ-irradiation and UVC light did not affect the interactions between GFP-MAP7 with endogenous BRCA1 and RAD50 or between mCherry-MAP7D1 with endogenous BRCA1 (Figure 5A-C).

**Figure 4.**
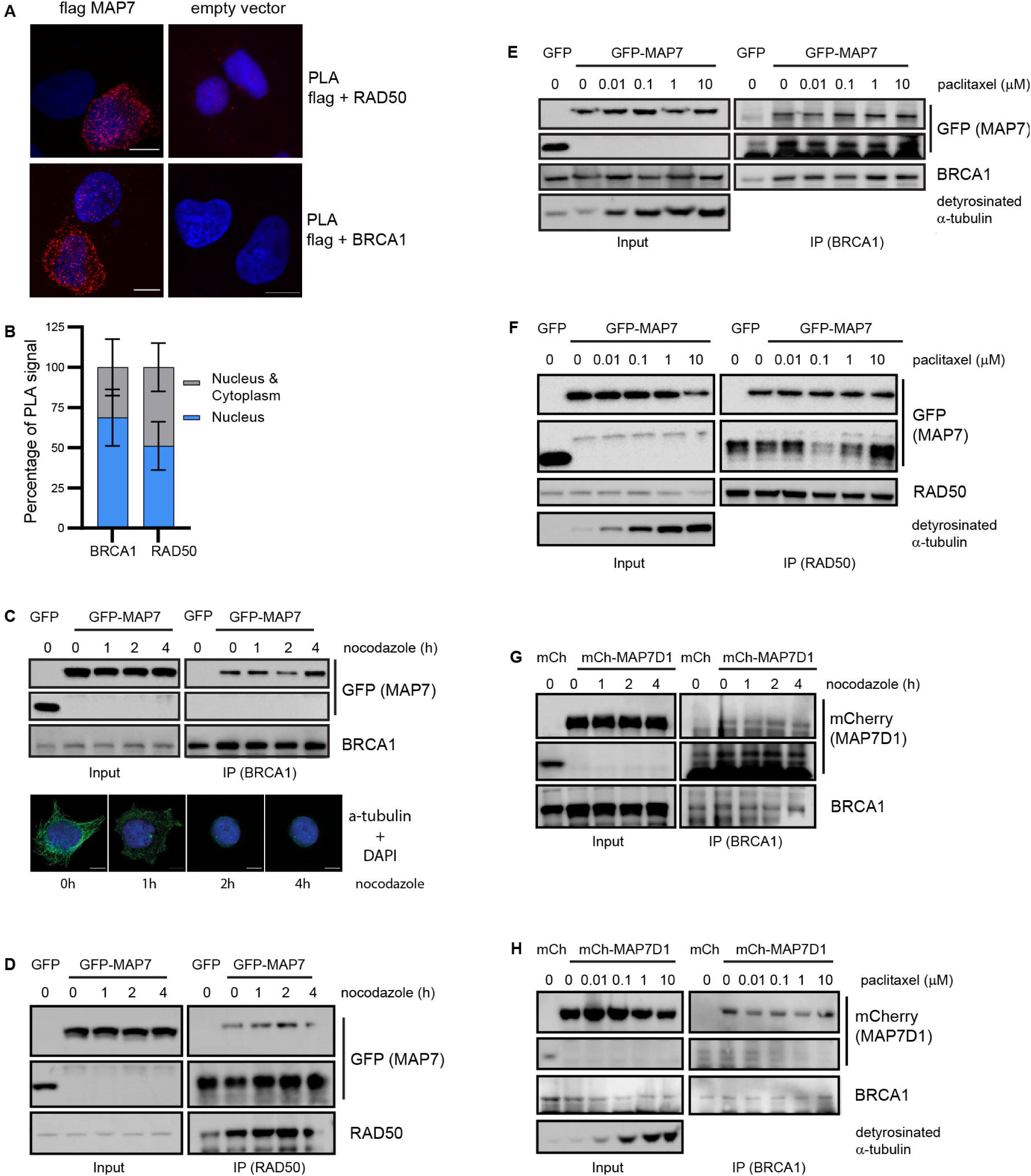
The interactions between MAP7/MAP7D1 and DDR proteins are not altered by nocodazole or paclitaxel. (A) Proximity ligation assay with endogenous RAD50 and Flag-MAP7 (top left) or empty Flag vector (top right) and with endogenous BRCA1 and Flag-MAP7 (bottom left) or empty Flag vector (bottom right) in U2OS cells. (B) Quantification of PLA staining showing the percentage of cells containing the PLA signal both in the cytoplasm and in the nucleus or only in the nucleus. (C) Top: immuno-precipitation of GFP-MAP7 with endogenous BRCA1 from HEK293T whole cell extracts from cells treated with 0.1 ng/ml nocodazole at different time points. Bottom: representative immuno-fluorescent images of microtubules in HEK293T treated with nocodazole at different time points. (D) Immuno-precipitation of GFP-MAP7 with endogenous RAD50 from HEK293T whole cell extracts from cells treated with 0.1 ng/ml nocodazole at different time points. (E) Immuno-precipitation of GFP-MAP7 with endogenous BRCA1 from HEK293T whole cell extracts from cells treated with increasing doses of paclitaxel (0-10 µM). Detyronisated α-tubulin is used as indicator of paclitaxel treatment. (F) Immuno-precipitation of GFP-MAP7 with endogenous RAD50 from HEK293T whole cell extracts from cells treated with increasing doses of paclitaxel (0-10 µM). (G) Immuno-precipitation of mCherry-MAP7D1 with endogenous BRCA1 from HEK293T whole cell extracts from cells treated with 0.1 ng/ml nocodazole at different time points. (H) Immuno-precipitation of mCherry-MAP7 with endogenous BRCA1 from HEK293T whole cell extracts from cells treated with increasing doses of paclitaxel (0-10 µM).

**Figure 5.**
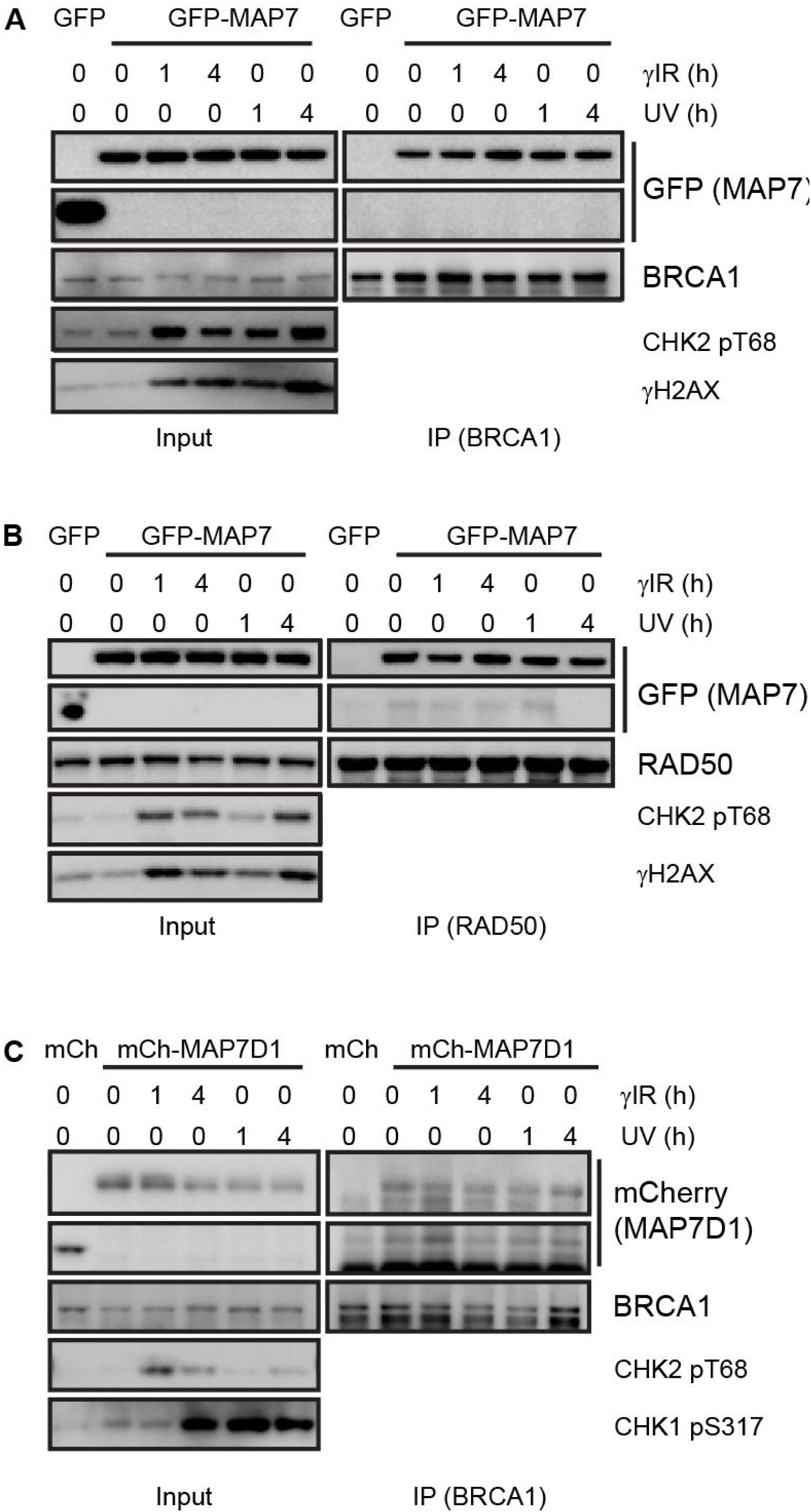
The interactions between MAP7/MAP7D1 and DDR proteins are not altered by DNA damage. (A) Immuno-precipitation of GFP-MAP7 with endogenous BRCA1 from HEK293T whole cell extracts from cells collected at 1 and 4 h after 10 Gy of γ-irradiation and 10 J/m^2^ UVC. CHK2 pT68 and γH2AX staining are used as indicators of irradiation treatment. (B) Immuno-precipitation of GFP-MAP7 with endogenous RAD50 from HEK293T whole cell extracts from cells collected at 1 and 4 h after 10 Gy of γ-irradiation and 10 J/m UVC. CHK2 pT68 and γH2AX staining are used as indicators of irradiation. (C) Immuno-precipitation of mCherry-MAP7D1 with endogenous BRCA1 from HEK293T whole cell extracts from cells collected at 1 and 4 h after 10 Gy of γ-irradiation and 10 J/m UVC. CHK2 pT68 and CHK1 pS317 staining are used as indicators of irradiation treatment.

### Downregulation of MAP7 and MAP7D1 leads to defects in DNA repair

Since microtubules and kinesins have been linked to the repair of DSBs^12, 14^, we sought to determine if MAP7 and MAP7D1 affect the repair of DNA damage caused by γ-irradiation. We used RPE1, MCF7 and A549 cell lines as they have wild-type p53 but differ in the ratio of expressed MAP7 and MAP7D1 (Figure 6A, B and Figure S6A). MAP7 and MAP7D1 were downregulated using three different siRNAs for each mRNA (Figure S6C). As siRNA 2 was less efficient in the downregulation of both MAP7 and MAP7D1 than siRNA 1, we have used siRNA 1 (MAP7#1) and a mix of two siRNAs (MAP7#2+3) in most of our experiments as a control for siRNA side effects. We did not detected any significant difference in phosphorylation of γH2AX and CHK2 Tyr68 on western blot at different time points after γ-irradiation, indicating that the activation of DDR through ATM is not affected by the downregulation of MAP7 and MAP7D1 (Figure 6A, B). However, we observed higher levels of total p53 and p53 Ser15 phosphorylation after γ-irradiation in cells with downregulated MAP7 and MAP7D1 simultaneously (Figure. 6A-D; Figure S6A, B) but not separately (Figure 6E, F). These results suggest that simultaneous downregulation of MAP7 and MAP7D1 either directly affects the stability and phosphorylation of p53 or that it causes defects in DNA repair. Since several cytoskeleton-related proteins such as KIF18B, LaminA/C and Lamin B1 have been reported to affect 53BP1 recruitment to the site of the damage^14, 15, 36, 37^, we tested if the downregulation of MAP7 and MAP7D1 affects the number of 53BP1 foci after γ-irradiation. Interestingly, MAP7D1 (but not MAP7) knock-down led to significantly fewer 53BP1 foci after γ-irradiation in MCF7 and RPE1 cells (Figure 5G-I).

**Figure 6.**
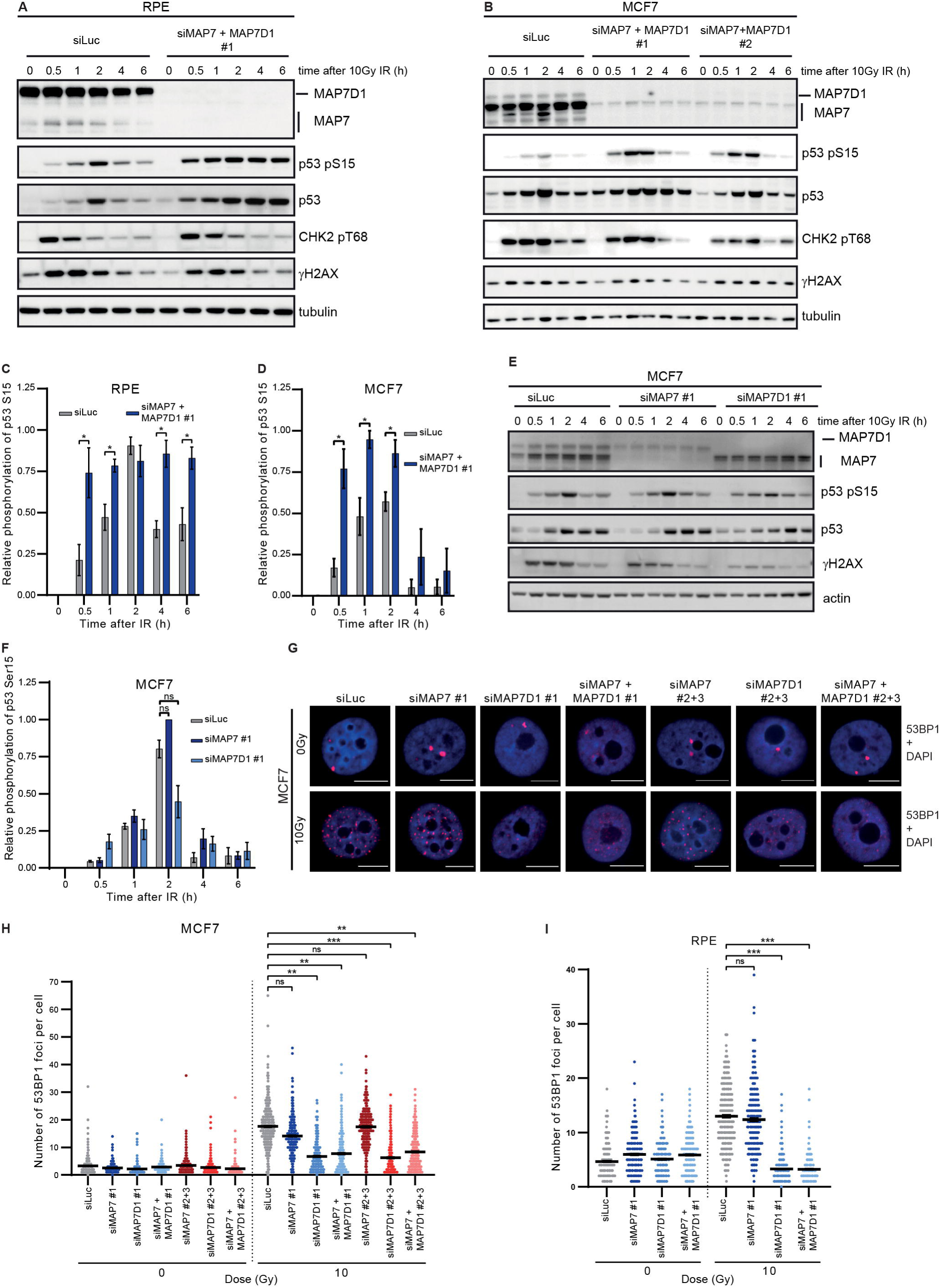
Downregulation of MAP7 and MAP7D1 leads to defects in DNA repair. (A) Western blot analysis of DNA damage response after 10 Gy of γ-irradiation in RPE1 cells treated with siRNA targeting luciferase or MAP7 and MAP7D1 at different time points. (B) Western blot analysis of DNA damage response after 10 Gy of γ-irradiation in MCF7 cells treated with siRNA targeting luciferase or MAP7 and MAP7D1 (two types of siRNA) at different time points. (C) Quantification of relative phosphorylation of p53 on Ser15 after γ-irradiation in RPE1 cells treated with siRNA targeting luciferase or MAP7 and MAP7D1. Signal intensity quantification was performed using ImageJ software. The p-Ser15 of p53 signal intensity was normalised against β-actin and expressed as relative phosphorylation of p53 at Ser15. Results are presented as mean ± SEM of values from three independent experiments. Statistical differences were determined from unpaired *t*-test: *p*<0.05, *. (D) Quantification of relative phosphorylation of p53 on Ser15 after γ irradiation in MCF7 cells treated with siRNA. Signal intensity quantification was performed using ImageJ software. The p-Ser15 of p53 signal intensity was normalised against β-actin and expressed as relative phosphorylation of p53 at Ser15. Results are presented as mean ± SEM of values from three independent experiments. Statistical differences were determined from unpaired *t*-test: *p*<0.05, *. (E) Western blot analysis of DNA damage response after 10 Gy of γ-irradiation in MCF7 cells treated with siRNA targeting luciferase, MAP7 or MAP7D1 at different time points. (F) Quantification of relative phosphorylation of p53 on Ser15 after gamma irradiation in MCF7 cells treated with siRNA. Signal intensity quantification was performed by ImageJ software. The p-Ser15 of p53 signal intensity was normalised against β-actin and expressed as relative phosphorylation of p53 at Ser15. Results are presented as mean ± SEM of values from three independent experiments. Statistical differences were determined from unpaired *t*-test: *p*<0.05, *, ns, not significant. (G) Immunofluorescence staining of 53BP1 foci in MCF7 cells untreated or treated with γ- irradiation (2 hours after irradiation). Scale bar, 10 µm. (H) Quantification of the immunofluorescence staining in MCF7 cells shown in (G). Data from three independent biological replicates are shown in the dot plot from which at least 300 cells were quantified. Results are presented as mean ± SEM of values. Statistical differences were determined from nested *t*-test: *p*<0.01, **; *p*<0.001, ***; ns, not significant. (I) Quantification of the immunofluorescence staining in RPE1 cells. Data from three independent biological replicates are shown in the dot plot from which at least 300 cells were quantified. Results are presented as mean ± SEM of values. Statistical differences were determined from nested *t*-test: *p*<0.001, ***; ns, not significant.

Given the difference in the 53BP1 recruitment in the cells treated with MAP7 and MAP7D1 siRNA, we investigated whether the downregulation of MAP7D1 causes specific cell cycle arrest. Indeed, using FACS analysis we observed that the knock-down of MAP7D1 leads to strong arrest in the G1 cell cycle phase in MCF7 and RPE1 cells (Figure 7A, B). We therefore speculated that MAP7 and MAP7D1 may play a role in DNA repair in the G1 cell cycle phase and examined if the knock-down of MAP7 in G1 arrested cells also leads to defects in 53BP1 recruitment to the site of the damage. We arrested MCF7 cells in G1 using the CDK4/6 inhibitor palbociclib, and treated them with 10Gy γ-irradiation. In these conditions, the down-regulation of both MAP7 and MAP7D1 led to fewer 53BP1 foci (Figure 7C). Consistently, alkaline comet assay experiments identified that the knock-down of MAP7 and MAP7D1 led to increased levels of DNA damage two hours after 8Gy of γ- irradiation in G1 arrested cells (Figure 7D, E). These results suggest that DNA repair is impaired in cells with downregulated MAP7 and MAP7D1 in the G1 cell cycle phase and that MAP7 and MAP7D1 affect the recruitment of 53BP1 to the site of the damage. We then wanted to see if the downregulation of MAP7 and MAP7D1 has a similar effect on the localization of BRCA1 and RAD50 to the chromatin. Since BRCA1 does not bind to the DNA damage foci in the G1 phase and since RAD50 staining was not possible to reliably interpret in our cells, we performed cell fractionation followed by the detection of chromatin-bound BRCA1 and RAD50 by western blot. Although the levels of BRCA1 in MCF7 G1 arrested cells were too low for detection, we were able to see that RAD50 was depleted from chromatin in cells treated with MAP7 or MAP7D1 siRNA both before and after 10Gy γ-irradiation (Figure 7F, G). Our data suggest that MAP7 and MAP7D1 affect the recruitment of the DDR proteins to chromatin and/or DNA double-strand breaks in the G1 cell cycle phase.

**Fig. 7.**
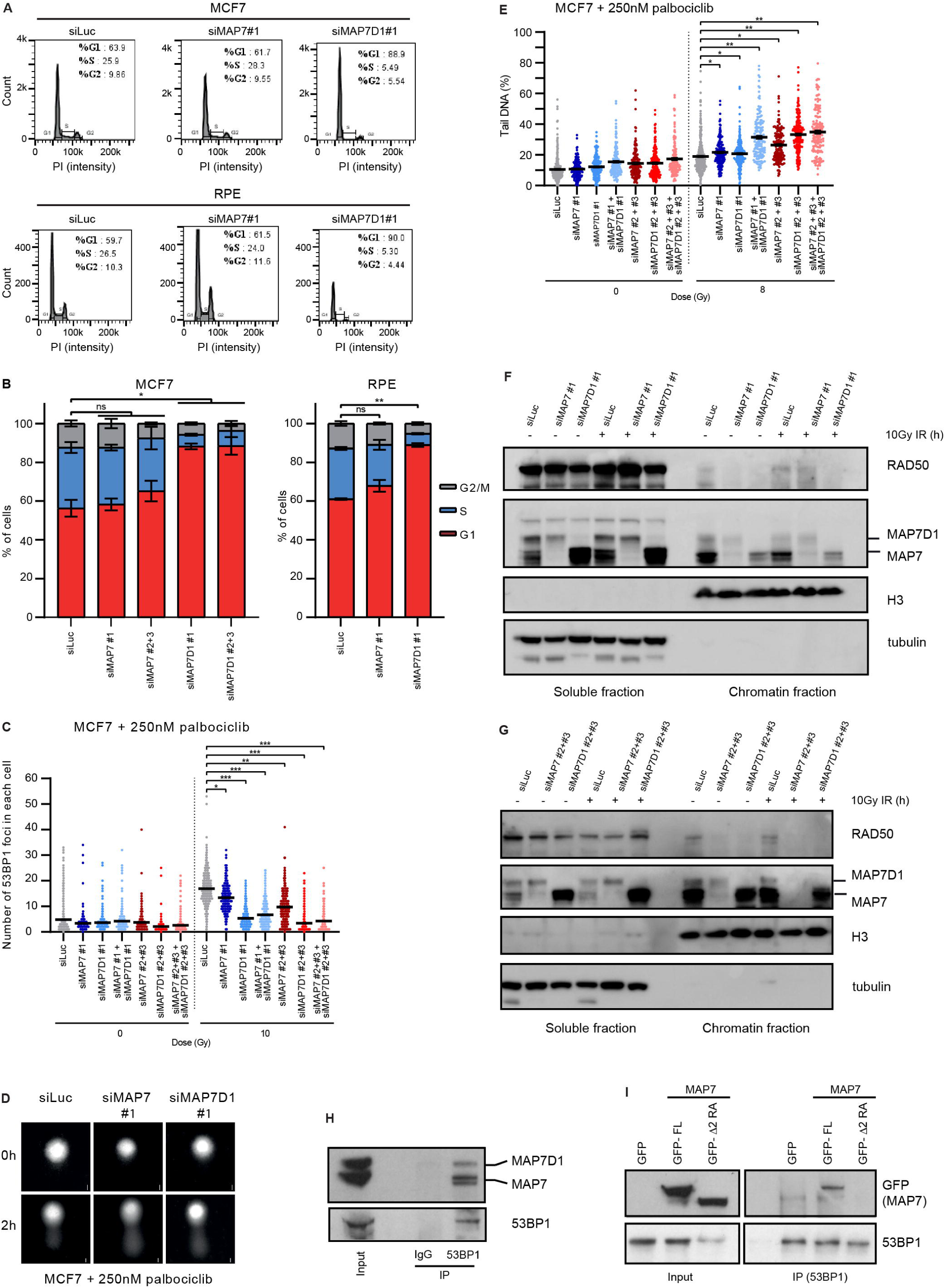
Knock-down of MAP7D1 leads to arrest in G1, while downregulation of MAP7 and MAP7D1 in G1 cells leads to defects in DNA repair. (A) Representative images of cell cycle analysis of MCF7 (top) and RPE1 cells (bottom) treated with siLuc, siMAP7 or siMAP7D1 siRNA. (B) Quantification of FACS cell cycle data from MCF7 (right) and RPE1 (left) cells. Data from three independent biological replicates are shown in the bar graph from which on average 30,000 cells for MCF7 and 10,000 cells for RPE1 were acquired. Results are presented as mean ± SEM of values. Statistical differences were determined from nested *t*-test: *p*<0.05, *; *p*<0.01, **, ns, not significant. (C) Quantification of the immunofluorescence staining of 53BP1 foci in MCF7 cells untreated or treated with 10Gy γ -irradiation. Cells were treated with siRNA targeting luciferase, MAP7 or MAP7D1 and arrested in the G1 cell cycle phase using 250 nM palbociclib. Data from three independent biological replicates are shown in the dot plot from which at least 300 cells were quantified. Results are presented as mean ± SEM of values. Statistical differences were determined from nested *t*-test: *p*<0.05, *; *p*<0.01, **; *p*<0.001, ***. (D) Representative images of alkaline comet assay with MCF7 cells arrested in G1 cell cycle phase using 250 nM palbociclib. The cells were downregulated with siRNA targeting luciferase, MAP7 or MAP7D1 and treated or untreated with 10 Gy of γ-irradiation. - Scale bar, 10µm. (E) Quantification of alkaline comet assay experiments with MCF7 cells from (D). Data from three independent biological replicates are shown in the dot plot from which at least 150 cells were quantified. Results are presented as mean ± SEM of values. Statistical differences were determined from nested *t*-test: *p*<0.05, *; *p*<0.01, **. (F) and (G) Western blot analysis of MCF7 cell fractionation. MCF7 cells were treated with siRNA and treated or untreated with 10 Gy of γ-irradiation and collected 1 h after the treatment. (H) Immuno-precipitation of endogenous MAP7 and MAP7D1 with 53BP1 from HEK293T whole cell extract (both MAP7 and MAP7D1 are recognised by MAP7 antibody). (I) Immuno-precipitation of full length MAP7 (GFP-FL) wild type and MAP7 mutated on Arg144/145 to Ala with deletion between aa151-300 (GFP- Δ 2) with endogenous 53BP1 from HEK293T whole cell extract.

Since MAP7 and MAP7D1 bind to DNA repair proteins including RAD50 and BRCA1, which are directly involved in DSB repair, we next investigated if MAP7 and MAP7D1 could affect 53BP1 recruitment through a direct interaction. Immunoprecipitation experiments revealed that endogenous 53BP1 interacts with endogenous MAP7 and MAP7D1 (Figure 7H). Further immunoprecipitation experiments revealed that the interaction between MAP7 and 53BP1 is (similarly to BRCA1, RAD50, MLH1 and XPC) mediated by the MAP7 region between aa151-300 (Figure 7I). These data imply that MAP7 and MAP7D1 may regulate recruitment of 53BP1 to the site of the damage through their direct interaction.

Overall, our findings describe for the first time an important function of MAP7 and MAP7D1 in cell cycle regulation and cellular response to DNA damage caused by γ-irradiation.

## Discussion

Microtubule-associated proteins (MAPs) regulate many important cellular processes such as cell motility, division and differentiation^17, 18^. Both MAP7 and MAP7D1 play roles in the recruitment and activation of kinesin1 to the microtubules^19–26^, are involved in cargo transport and in the β-catenin independent Wnt5a signalling^38^. High expression of MAP7 (also known as ensconsin or E-MAP-115) has been reported to be an adverse prognostic biomarker for cytogenetically normal acute myeloid leukaemia, stage II colon cancer, cervical cancer and metastatic endometrial cancer^39–41^. Additionally, high expression levels of MAP7D1 have been also recently shown to promote breast cancer proliferation and metastasis^42^.

Here we show that both MAP7 and MAP7D1 bind several DNA repair proteins including BRCA1, RAD50, MLH1, XPC and 53BP1. We have discovered that the direct interactions with RAD50 and BRCA1 are dependent on the CK2 phosphorylated sites within the DNA repair proteins. In addition, our data strongly suggest that the interactions between the DDR proteins and MAP7 are also bridged by other protein(s) and that these interactions are phosphorylation independent. The region of MAP7 required for the direct and indirect interactions spans between aa144-300. Interestingly, a similar but not identical region of MAP7 (aa89-246) has been shown to interact with DVL2, which is involved in β-catenin-independent Wnt signalling^38^. Although the structured regions of MAP7 and MAP7D1 (CC1 and CC2) share similar sequences, of the other regions of these proteins are predicted to be unstructured. Our results indicate that the deletion of a similar region of MAP7D1 does not lead to complete loss of interaction with the DDR proteins. Since only part of the deleted region contains the conserved CC1 domain and the majority is within the unstructured region, both proteins may interact with the DDR proteins through different regions. Our experiments revealed that the downregulation of MAP7D1 but not MAP7 in MCF7 and RPE1 cells leads to strong G1 arrest. Previously published data show that downregulation of MAP7 leads to arrest in G2 in SiHa cells. This cell line is HPV positive, thus suppression of the p53 pathway and of the G1 checkpoint is expected^28, 43^.

We have discovered that MAP7 (and possibly MAP7D1) is exclusively involved in G1 DNA repair as we only detected effects on 53BP1 foci in G1 arrested cells. Importantly, other cytoskeleton-related proteins such as KIF18B, KIF2C, Lamin A/C and Lamin B have also been shown to regulate the recruitment of 53BP1 to the site of the damage, however in contrast to MAP7 and MAP7D1, throughout the whole cell cycle^14, 15, 36, 37^. This strongly suggests that MAP7 and MAP7D1 affect 53BP1 recruitment independently on these factors.

In addition, DSB repair is also affected by motility of microtubules. The interaction between the microtubule network with the linker of nucleoskeleton and cytoskeleton (LINC) together with 53BP1 promotes DSB repair through increased mobility of DSBs by microtubules^11, 44^. At the same time, DNA damage itself promotes microtubule dynamics in G0/G1 cell cycle phase, which is regulated through DNA-PK and AKT signalling^12^. Although the microtubule motility directly affects DSB repair, it does not affect 53BP1 recruitment, indicating that the mechanism through which MAP7 and MAP7D1 are involved in DDR may be different.

Interestingly, we see a stronger phenotype for MAP7D1 downregulation in terms of cell cycle arrest and 53BP1 recruitment in both RPE1 and MCF7 cells, although MCF7 cells express much higher levels of MAP7 compared to MAP7D1. It is therefore possible that MAP7D1 plays a more distinct role in cell cycle regulation and DDR. This would be in line with the fact that MAP7D1 but not MAP7 was found to be one of the proteins important for DDR by a large study involving CRISPR-Cas9 screens with 27 genotoxic agents in p53 -/- RPE1 cells^45^. On the other hand, our data also a show similar effect of MAP7 and MAP7D1 downregulation on RAD50 binding to chromatin in G1 cells in non-damage and damage conditions, indicating that the function of MAP7 and MAP7D1 may vary with different interacting DDR proteins.

Taken together, our data show for the first time that MAP7D1 plays a role in the regulation of the cell cycle, that MAP7 and MAP7D1 bind several DNA repair proteins and that they are involved in DDR in the G1 phase through the regulation of 53BP1 recruitment to the site of the damage and RAD50 recruitment to chromatin. In light of recent discoveries that motor activities of kinesins KIF2C and KIF18B are important for efficient DSB repair and that cytoplasmic microtubules are required for DNA repair through the regulation of the mobility of DSBs, our data may contribute to the hypothesis that (similarly to the role of actin filaments in DSB repair) there could be a subset of short and transient microtubules in the nucleus (so far undetected in mammalian cells) that can facilitate DNA repair. Alternatively, MAP7 and MAP7D1 may be part of several factors that are involved in microtubule assembly/disassembly, which may be connecting the cytoplasmic and nuclear cytoskeleton, and which may mediate the repair of damaged DNA. Future work is required to determine the precise mechanism by which MAP7 and MAP7D1 are involved in the DDR and cell cycle regulation and to elucidate if the microtubule associated proteins may be linking the cytoskeleton-related proteins with the DDR machinery.

## Limitations of the study

Although our study identifies the effect of the downregulation of MAP7 and MAP7D1 on the recruitment of 53BP1 to the site of damage and RAD50 to the chromatin, we have not been able to determine the effect of MAP7 and MAP7D1 on the recruitment of BRCA1. Since the effects were observed in the G1 phase, when BRCA1 levels are low and it is not recruited to the site of the damage, we have not been able to establish an assay that would assess the function of the MAP7/MAP7D1 interaction with BRCA1. It was also not in the scope of this work to identify the mechanism by which MAP7/MAP7D1 affect the recruitment of RAD50 and 53BP1.

## Materials and Methods

### Cell culture, siRNA and drug treatment

HEK293T, U2OS, MCF7, RPE1 and A549 cells (ATCC *via* The Francis Crick Institute Cell Services) were maintained as an adherent monolayer in DMEM media containing 10% FBS and 1% penicillin/streptomycin at 37°C in a humidified atmosphere of 5% carbon dioxide. For siRNA transfections, 6µl of Lipofectamine RNAiMAX Reagent (Invitrogen) was diluted in 100µl of DMEM and separately, 400nM of each siRNA was diluted in 100µl of DMEM. After incubation for 5 min at room temperature, they were combined and incubated for 20 min at room temperature. The mixture was added to cells with 800µl of DMEM + 10% FBS and incubated for 5 h before media was refreshed. For high knockdown efficiency, siRNA transfection was performed 24 h and 72 h after seeding.

Cells were treated with 10Gy γ-IR, 0.1ng/ml nocodazole (Sigma-Aldrich), 0.01µM-10µM paclitaxel (TOKU-E), 250nM palbociclib (Selleck), 100 or 500 units of recombinant CK2 (NEB), 5µM or 15µM silmitaserib (CK2 inhibitor, Cayman Chemical) and 10J/m^2^ UVC.

### Mass spectrometry analysis and protein identification

Immunoprecipitated protein complex beads were digested into peptides using 500ng sequencing grade trypsin (Thermo Fisher Scientific) and incubated overnight at 37°C on a shaker. Peptides were then desalted using C18+carbon top tips (Glygen Corporation, TT2MC18.96) and eluted with 70% acetonitrile (ACN) with 0.1% formic acid.

Dried peptides were dissolved in 0.1% TFA and analysed by nanoflow ultimate 3000 RSL nano instrument that was coupled on-line to a Q Exactive plus mass spectrometer (Thermo Fisher Scientific). Gradient elution was from 3% to 28% buffer B (0.1% formic acid in ACN) in 120 mins at a flow rate of 250nL/min with buffer A (0.1% formic acid in water) being used to balance the mobile phase. The mass spectrometer was controlled by Xcalibur software (version 4.0) and operated in the positive mode. The spray voltage was 1.9kV and the capillary temperature was set to 255°C. The Q-Exactive plus was operated in data dependent mode with one survey MS scan followed by 15 MS/MS scans. The full scans were acquired in the mass analyser at 375-1500m/z with the resolution of 70,000, and the MS/MS scans were obtained with a resolution of 17,500.

MS raw files were converted into Mascot Generic Format using Mascot Distiller (version 2.5.1) and searched against the SwissProt database (release December 2015) restricted to human entries using the Mascot search daemon (version 2.5.0) with a FDR of ∼1% and restricted to the human entries. Allowed mass windows were 10ppm and 25mmu for parent and fragment mass to charge values, respectively. Variable modifications included in searches were oxidation of methionine, pyro-glu (N-term) and phosphorylation of serine, threonine and tyrosine. The mascot result (DAT) files were extracted into excel files for further normalisation and statistical analysis. All downstream data analysis was performed by Perseus (version 1.5.5.3)^46^. Normalized MS2 intensities were converted to Log2 scale. Reverse (decoy) hits, potential contaminants, and proteins identified only by modified peptides were filtered out. Ratio values were then median subtracted. Volcano plots of the MS2 intensities ratio values were also generated by Perseus. Benjamini-Hochberg cut-off of 5% with an s0 of (0.1) was used to select significant binders.

### Plasmids

MAP7 cDNA was purchased from Dharmacon and cloned to pDONR221 (Invitrogen), from which it was cloned by Gateway LR reaction to pDEST-FTF/FRT/TO^47^ or pDEST53 plasmid (Invitrogen) for mammalian expression. The GFP-BRCA1 expression plasmid was a gift from the Solomon lab^48^, the Flag-RAD50 expression plasmid was a gift from the Staples lab^49^ and the mCherry MAP7D1 was a gift from the Akhmanova lab^21^. The R144/145A and the ΔN1 and ΔN2 deletion mutations of MAP7, the Δ MAP7D1 deletion mutation, T690A mutation of RAD50 and S1336A mutation of BRCA1 were introduced using the Q5 Site-directed mutagenesis kit (NEB). The MAP7 truncation mutations were created by PCR cloning to Gateway p221 pDONR221 (Invitrogen), from which it was cloned by Gateway LR reaction to pDEST-FTF/FRT/TO^47^ or pDEST53 plasmid (Invitrogen) for mammalian expression.

### Protein extracts, immunoprecipitation and pull-down assays

Protein extracts were prepared from cells by lysing in immunoprecipitation (IP) buffer (50 mM Tris-HCl (pH 7.5), 150 mM NaCl, 1% Triton X-100, 1 mM EDTA, 2.5 mM EGTA, 10% (v/v) glycerol supplemented with cOmplete EDTA-free protease inhibitor (Sigma), and sonicated 3 × 10 s. The extracts were cleared by centrifugation at 10000 g for 15 min at 4°C.

For immunoprecipitations and pull-downs, 2.5 mg of the whole cell extract was incubated for 2 h at 4°C with 15 μl of M2-anti Flag agarose (Sigma-Aldrich), GFP-trap agarose (Chromo Tek) or Protein G agarose (Cell Signaling) bound with primary antibody. Beads were pelleted and washed three times in 20× bed volume of the lysis buffer. For immuno-precipitation, the bound protein was eluted by boiling in 2× LSB buffer (100 mM Tris (pH 6.8), 200 mM dithiothreitol (DTT), 4% sodium dodecyl sulfate (SDS), 0,2% bromophenol blue, 20% (v/v) glycerol) for 5 min. For pull-down, the bound protein was eluted by incubation with 30 µl of Flag peptide (according to manufacturer instructions for M2 agarose beads). Inputs represent ∼1/20th of the extract used for the immunoprecipitation. For clarity, some immuno-precipitates shown in the figures are from longer or shorter exposures than that shown for their corresponding inputs. Prior to λ- phosphatase treatment, Flag peptide elution or incubation with Flag-tagged eluted proteins, the beads (protein G, GFP-trap or M2 agarose) were successively washed in lysis buffer and wash buffer containing high salt (50mM Tris-HCl pH 8.0, 0.5-4M NaCl, 1% Triton X-100, 1 mM DTT, 1 mM EDTA). Treatment with λ-phosphatase (NEB) was performed as described previously^50^.

For the pull-down assay, 30 μl of the GFP-trap beads with bound GFP-tagged proteins and 60 μl of eluted Flag-tagged proteins were used per reaction. The GFP-trap beads were incubated with the eluted Flag-tagged proteins for 1 h at 4°C, washed three times in 20× bed volume of the lysis buffer and proteins were eluted from the GFP-trap agarose beads by boiling in 2× LSB buffer for 5 min.

The peptide pull-down was performed as described previously^31^ and carried out using biotinylated peptides synthesised by The Francis Crick protein chemistry facility

### Kinase assay

Beads with bound Flag-RAD50 or GFP-BRCA1 were washed with 5M NaCl to eliminate contamination with protein kinases and incubated with 100 or 500 U of CK2 holoenzyme complex (α2/β; NEB) in 30 μl of kinase buffer (20 mM Tris-HCl (pH 8.0); 50_mM KCl; 10 mM MgCl_2_, 33 μM ATP) and incubated for 30 min at 37 °C. Reaction was stopped by the addition of 4 × Laemmli buffer (200 mM Tris (pH 6.8), 400 mM dithiothreitol (DTT), 8% sodium dodecyl sulfate (SDS), 0.4% bromophenol blue, 40% (v/v) glycerol).

### Proximity ligation assay

U2OS cells grown on coverslips were transfected with Flag-RAD50 or empty Flag-plasmid. Cells were permeabilized with 0.2% Triton X-100 for 5 min at room temperature and proximity ligation assay (PLA) was performed using Flag, MAP7 antibodies and Duolink reagent (Sigma-Aldrich) according to the manufacturer’s protocol. For PLA quantification Z-stack images were taken for each condition with the LSM710 microscope at x63 objective lens. PLA signal was used to define the axis and 10 z-stack slices were acquired per image, stacks were pulled together, and merged with DAPI for the analysis.

### Immunofluorescence

Cells grown on glass coverslips were treated with siRNA. For G1-phase arrest, cells were treated with 250nM palbociclib for 24 h following the second round of siRNA treatment. Cells were treated with γ-IR then fixed after 2 h in 4% PFA in PBS for 10 min at room temperature and washed three times with PBS. Cells were permeabilised with PBS + 0.5% TX-100 for 10 min and incubated with the primary antibody in DMEM + 10% FBS for 1 hr at room temperature. The coverslips were washed three times with PBS and incubated with the secondary antibody in DMEM + 10% FBS 1 h at room temperature in the dark. Coverslips were washed with PBS three times and incubated in PBS + 0.2µg/ml DAPI for 2 min at room temperature. Coverslips were washed in distilled water, air-dried then mounted onto glass slides using ProLong Gold Antifade Mountant (Invitrogen). Images were acquired using Zeiss LSM 710 Confocal Microscope using a 63× objective lens. 53BP1 foci quantification was performed using ImageJ. Cell nuclei were defined by thresholding the DAPI signal and the regions of interest were saved to the ImageJ ROI manager. The 53BP1 signal was smoothed with the ‘Smooth’ function and the noise tolerance was adjusted using the ‘Find maxima’ command to identify the 53BP1 foci. Foci for each pre-selected ROI were automatically counted and results were presented as the sum of all pixels in the region on the ROI (RawIntDen). The pixel value is 255 so the sum of all pixels was divided by 255 to acquire the number of 53BP1 foci in the defined nuclear signal.

### FACS analysis

siRNA treated cells were harvested, washed with cold PBS and fixed by adding ice-cold 70% ethanol dropwise while vortexing. Cells were then incubated on ice for 30 min. Following that, the fixed cells were washed twice with PBS and treated with 50 µg/ml RNAase A for 30 min at 37 ° C. Finally, cells were stained with 20µg/ml PI on ice for 30 min. Flow Cytometry was performed on a BD LSR Fortessa cell analyser. On average 30,000 cells for MCF7 and 10,000 cells for RPE1 have been acquired. Data were analysed with Flow Jo software using the Cell Cycle Analysis Tool.

### Alkaline comet assay

The alkaline comet assay was performed using the Reagent Kit for Single Cell Gel Electrophoresis Assay (Trivegen) with adaptations to the manufacturer’s protocol. Cells were seeded and treated with siRNA followed by 250nM palbociclib for 24 h. Cells were collected 2 h h after 8 Gy γ-irradiation and suspended at 1× 10 cells/ml. The cells were mixed with LMAgarose and pipetted onto the slide. Slides were incubated at 4°C in the dark for 1 hr then incubated in Lysis Solution overnight at 4°C in the dark. Slides were incubated in Alkaline Unwinding Solution for 1 hr at 4°C in the dark. For electrophoresis, slides were placed in a horizontal electrophoresis tank containing Alkaline Electrophoresis Solution and electrophoresis was carried out for 25 min at 30V and around 300mA at room temperature. The slides were washed twice for 5 min in distilled water then fixed in 70% ethanol for 5 min, dried at 37°C and stored at 4°C. For staining of DNA, slides were immersed in SYBR Gold in TE Buffer (10mM Tris-HCl pH 7.5, 1mM EDTA) for 30 min at room temperature in the dark then washed with water and dried at 37°C. Imaging was gathered using Zeiss LSM 710 Confocal Microscope 10× objective lens. Semi-automated image analysis was used to analyse 50 individual comets per gel to calculate the percentage of tail DNA (https://www.med.unc.edu/microscopy/resources/imagej-plugins-and-macros/comet-assay/).

### Cellular fractionation

siRNA treated cells were incubated with 250nM palbociclib for 24 h then were collected 1 h after 10 Gy irradiation. Cell pellets were resuspended in CSK Buffer (10mM PIPES KOH pH 6.8, 100 mM NaCl, 300 mM sucrose, 1.5 mM MgCl_2_, 5 mM EDTA, 0.5% Triton X-100, 1× protease and phosphatase inhibitor, Sigma) and incubated on ice for 10 min. Cells were centrifuged at 3000 rpm for 3 min at 4°C and the supernatant was collected as the soluble fraction. The pellet was washed once more with CSK buffer and centrifuged at 3000 rpm for 3 min at 4°C. The supernatant was discarded and the pellet was collected as the chromatin fraction.

### Antibodies

The anti-phospho BRCA1 antibody was raised and purified by GenScript against phosphorylated peptide CQSESQGVGL{pSer}DKEL.

### Statistical analysis

All experiments were performed three times independently. Statistical analysis was performed using GraphPad Prism 9 software. Statistical details for the experiments are provided in the figure legends. *p*<0.05 was considered to be significant and classified by asterisks: *p*<0.05 (*), *p*<0.01 (**), *p*<0.001 (***) and *p*<0.0001 (****).

## Acknowledgements

This work was supported by the Wellcome Trust (200462/Z/16/Z). AD was supported by QMUL incentivisation award. We thank Christopher J Staples for providing us with the Flag-RAD50 expression plasmid, the Ellen Solomon lab for providing us with the GFP- BRCA1 expression plasmid and Anna Akhmanova lab for providing us with the mCherry- MAP7D1 plasmid. We thank Faraz Mardakheh for advice on the proteomic data analysis and visualisation. We are grateful to Roberto Bellelli and Libor Macurek for critical reading.

## Author Contributions

AD was responsible for planning and carrying out experimental work, data analysis and writing the report. MO was responsible for planning and carrying out experimental work, data analysis and writing the report. WYT carried out experimental work. VR carried out the work required for quantitative proteomics and its analysis. PRC provided feedback on the proteomics experiment planning and data analysis. SAM provided feedback on the experimental work and the report. ZH was responsible for planning and carrying out experimental work and writing the report.

## Data Availability Statement

Further information and requests for resources and reagents should be directed to and will be fulfilled by the Lead Contact Zuzana Horejsi. Plasmids and cell lines used in this work will be available upon request. All mass spectrometry raw files and search results reported in this paper have been deposited at the ProteomeXchange Consortium via the PRIDE partner repository with the dataset identifier PXD033715 and 10.6019/PXD033715.

---

**Figure S2:**
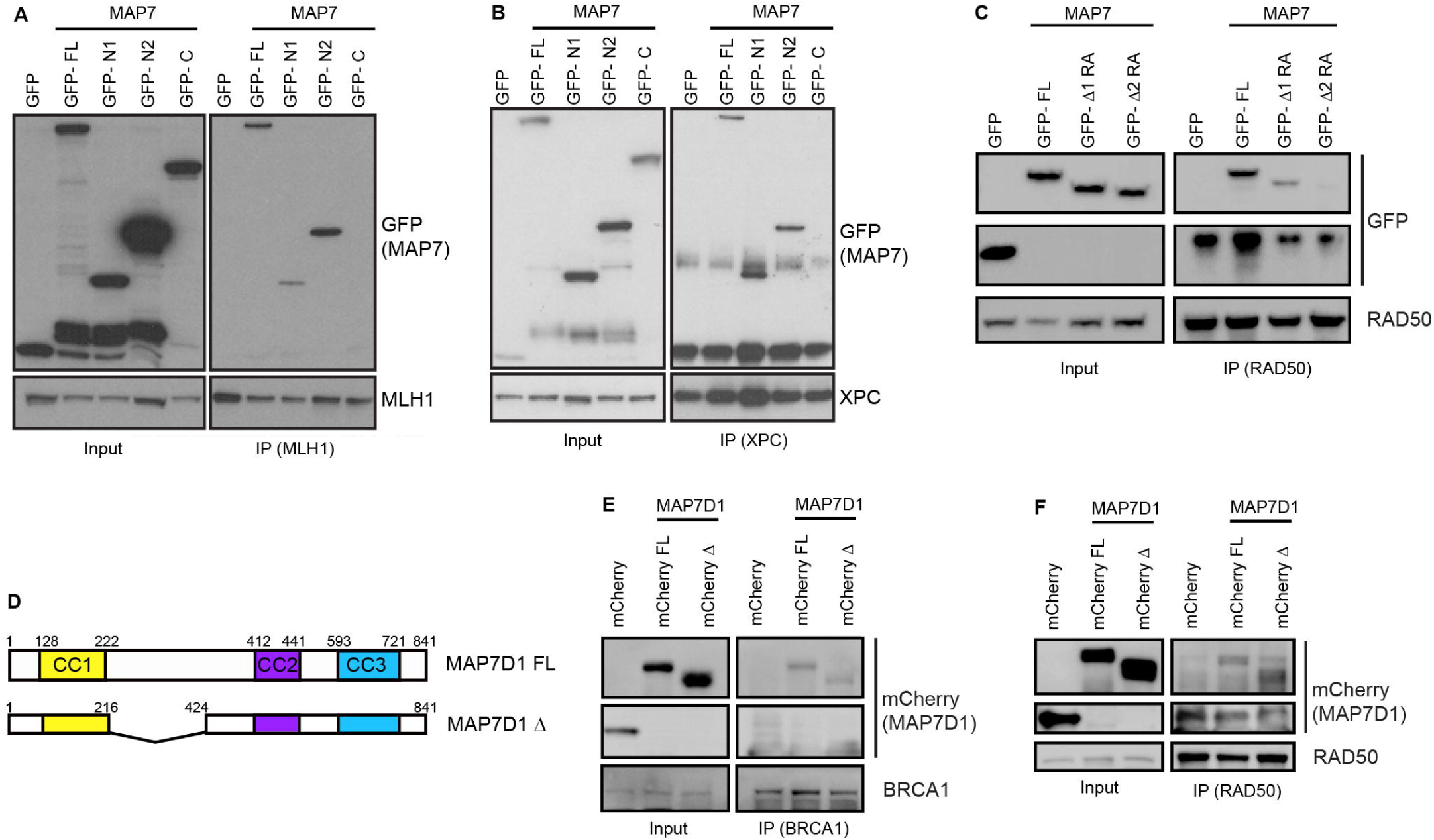
Interaction between MAP7 and DDR proteins MLH1 and XPC and domain mapping of MAP7D1. (A) Domain mapping by immuno-precipitation of full-length GFP-MAP7 (GFP-FL) and MAP7 truncation constructs by endogenous MLH1 from HEK293 extract overexpressing GFP-MAP7 constructs. (B) Domain mapping by immuno-precipitation of full-length GFP- MAP7 (GFP-FL) and GFP-MAP7 truncation constructs by endogenous XPC from HEK293 extract overexpressing GFP-MAP7 constructs. (C) Immuno-precipitation of GFP-MAP7 constructs mutated on Arg144/145 to Ala with deletion between aa151-270 (GFP-⊗N1 RA) and 151-300 (GFP-⊗N2 RA) with endogenous RAD50. (D) Schematic representation of full-length MAP7D1 and MAP7D1 deletion construct used in immunoprecipitations. (E) Immuno-precipitation of mCherry-tagged MAP7D1 wt and deletion mutation by endogenous BRCA1 from HEK293T extract overexpressing mCherry constructs. (F) Immuno-precipitation of mCherry-tagged MAP7D1 wt and deletion mutation by endogenous RAD50 from HEK293T extract overexpressing mCherry constructs.

**Figure S6.**
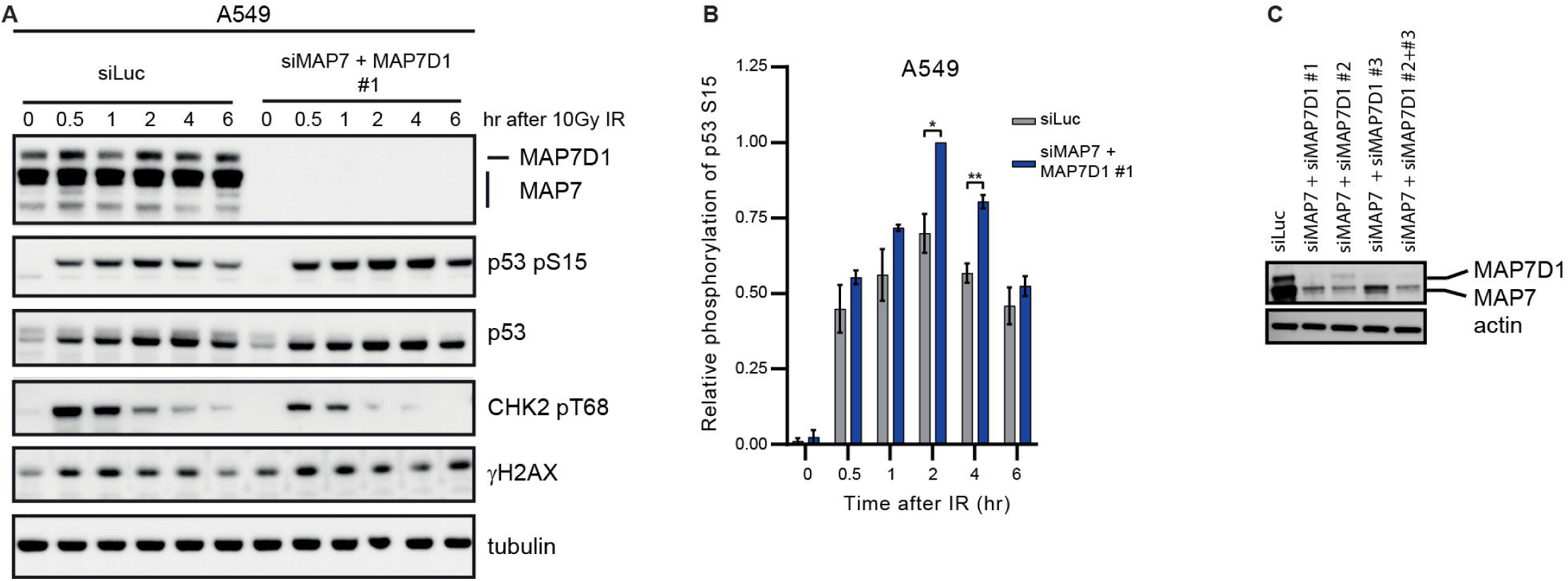
Downregulation of MAP7 and MAP7D1 leads to increased p53 S15 phosphorylation in A549 cells. (A) Western blot analysis of DNA damage response after 10Gy of gamma-irradiation in A549 cells treated with siRNA targeting luciferase or MAP7 and MAP7D1 at different timepoints. (B) Quantification of relative phosphorylation of p53 on Ser15 after γ-irradiation in A549 cells treated with siRNA targeting luciferase or MAP7 and MAP7D1. Signal intensity quantification was performed by ImageJ software. The p-Ser15 of p53 signal intensity was normalised against β-actin and expressed as relative phosphorylation of p53 at Ser15. Results are presented as mean ± SEM of values from three independent experiments. Statistical differences were determined from unpaired *t*-test: *p*<0.05, * *p*<0.01, **. (C) Western blot analysis of MAP7 and MAP7D1 siRNA treatment in MCF7 cell line.

